# NRF2 regulates a lipofuscin-like granule cycle through autophagy-dependent clearance in retinal pigment epithelium

**DOI:** 10.64898/2026.11.16.766427

**Authors:** Margarida L. Pedro, Shuvajit Rakshit, Mafalda Lopes-da-Silva, Leonor Devesa, Eduardo C. Ibáñez, Pedro Antas, António Cuadrado, Sandra Tenreiro, Ana S. Falcão, Miguel C. Seabra

**Affiliations:** iNOVA4Health, NOVA Medical School|Faculdade de Ciências Médicas, NMS|FCM, Universidade Nova de Lisboa, Lisboa, Portugal; Champalimaud Research, Champalimaud Foundation, Lisbon, Portugal, Lisboa, Portugal; Centro de Investigación Biomédica en Red Sobre Enfermedades Neurodegenerativas (CIBERNED), Instituto de Investigación Sanitaria La Paz (IdiPaz), Department of Biochemistry and Instituto de Investigaciones Biomédicas Alberto Sols UAM-CSIC, Faculty of Medicine, Autonomous University of Madrid, Madrid, Spain

**Keywords:** age-related macular degeneration, retinal pigment epithelium, NRF2, dimethyl fumarate, autophagy, lysosome dysfunction, lipofuscin

## Abstract

Lysosomal dysfunction in retinal pigment epithelial (RPE) cells is one pathological aspect of retinal degenerative disorders, which contribute to the buildup of autofluorescent undegraded materials such as lipofuscin. A comprehensive understanding of lipofuscin biology remains a critical gap in the field. Here, we explore the cell biology of lipofuscin-like autofluorescent granules (AFGs) within RPE cells and the potential of nuclear factor erythroid 2-related factor 2 (NRF2) activation to promote its clearance in human RPE cells. Under steady-state conditions we find that photoreceptor outer segments (POS) phagocytosis by the RPE is sufficient to trigger NRF2 activation, suggesting a feedback mechanism that links phagocytic activity with cytoprotective stress responses. RPE fed with excess POS accumulate lipofuscin-like autofluorescent granules (AFG) within hours, a measure of lysosomal dysfunction. These AFGs gradually loose connection with the endocytic system and remain as “storage granules” within RPE. Pre-treatment or delayed pharmacological activation of NRF2 with dimethyl fumarate (DMF) significantly decrease accumulation of AFGs, suggesting improved clearance mechanisms. Genetic or pharmacological inhibition of autophagy totally abolish these effects. Similarly, compromised autophagy also abolished the clearance effects observed with inhibition of the mTOR pathway. Collectively, these results suggest a 3-step lipofuscin cycle within the RPE (formation, storage and clearance), provide mechanistic evidence that autophagy is an essential process for the clearance of AFGs and that NRF2-activation and other stimulators of autophagy may represent novel therapeutic strategies to restore RPE homeostasis and mitigate retinal degeneration.

## Introduction

The retinal pigment epithelium (RPE) plays a critical role in maintaining photoreceptor health and visual function, in part through the daily phagocytosis of photoreceptor outer segments (POS). This high degradative demand renders RPE cells particularly reliant on efficient phagocytic pathways and lysosome function. Dysregulation of these pathways contributes to the accumulation of undegradable materials, such as lipofuscin [1]. Although lipofuscin accumulation is a recognized feature of normal aging, its excessive buildup is associated to pathological changes in the retina, like impaired lysosomal cargo degradation and autophagic flux, increased oxidative stress, inflammation, and cell death [2–4]. These changes can compromise the supportive role of the RPE and eventually lead to photoreceptor degeneration. Conditions such as Stargardts disease and age-related macular degeneration (AMD) have been postulated to involve lipofuscin-related pathology, which suggests that accumulation of this material is not merely a passive marker of aging, but may actively contribute to disease processes mainly due to lipid peroxidation [5,6].

In RPE cells, lysosomal homeostasis is essential for maintaining the capacity to process the continuous influx of photoreceptor outer segments (POS) and intracellular waste. Following phagocytosis, POS-containing phagosomes mature and interact with lysosomes, where their contents are degraded and cellular components are recycled. When degradation is incomplete, residual material can accumulate as autofluorescent granules (AFGs), which may progressively become less accessible to the endolysosomal pathway and persist as intracellular storage structures [7]. Macroautophagy provides an additional mechanism for maintaining cellular proteostasis by sequestering intracellular material within autophagosomes and delivering it to lysosomes for degradation. This process therefore depends not only on autophagosome formation but also on the availability and functionality of lysosomes. Lipofuscin precursor material is believed to enter RPE cells through phagocytosis of POS, generating phagosomes that subsequently fuse with lysosomes. Efficient phagosome processing relies on LC3-associated phagocytosis (LAP), in which LC3 is recruited to single-membrane phagosomes, promoting phagosome-lysosome fusion and degradation of POS [8–11]. Conversely, the role of canonical macroautophagy in POS phagocytosis is less clear. Given the role of macroautophagy in clearing mal-functioning organelles and protein aggregates, impairment of this process could potentially compromise POS processing, favouring the accumulation of incompletely degraded material that contribute to lipofuscin formation.

Understanding mechanisms of lipofuscin formation and degradation is therefore crucial. First, it may illuminate why some individuals experience accelerated retinal degeneration while others maintain retinal health despite aging. Second, identifying key regulators of lipofuscin homeostasis could inform the development of therapies aimed at modulating lysosomal function, enhancing autophagic flux, or preventing the formation of toxic bisretinoids, such as A2E (a major component of lipofuscin) [12]. Third, insights into lipofuscin dynamics could help refine current diagnostic tools, such as fundus autofluorescence (AF) imaging, which detects lipofuscin *in vivo* and is widely used to monitor disease progression in retinal disorders. Despite its central role in retinal aging and disease, effective interventions targeting lipofuscin accumulation and clearance remain limited. Unlike other cellular waste products, lipofuscin is resistant to enzymatic degradation and cannot be easily removed by exocytosis or lysosomal turnover. As a result, it tends to accumulate chronically within post-mitotic cells such as the RPE [1–5]. Recent research began to explore strategies to reduce lipofuscin burden, including the use of small molecules to inhibit A2E formation, gene therapies targeting visual cycle enzymes, and agents that stimulate lysosomal activity [6,13–15]We have recently shown that viral or pharmacological approaches activating the transcription factor EB (TFEB) pathway in the RPE could improve POS clearance and hence reduce AFG load [16]. Advancing our knowledge on how RPE cells cope with lipofuscin accumulation, and more importantly, how they might clear or neutralize it, holds great promise for preventing or slowing retinal degeneration.

Nuclear factor erythroid 2-related factor 2 (NRF2) is a transcription factor that plays a crucial role in regulating the cellular defence mechanisms. Its action on a wide range of physiological processes suggests NRF2 as a promising target for the development of novel treatments, including neurodegenerative, autoimmune, metabolic, cardiovascular diseases, and cancer [17–21]. NRF2 significance in AMD has been increasingly recognized: firstly, NRF2 signalling is impaired in RPE in aged mice and its deficiency leads to RPE damage, namely drusen-like deposits, increased AF and sub-RPE deposition of inflammatory proteins, resembling dry AMD, one of the advanced forms of the disease and the most common one [22]; secondly, a single-nucleotide polymorphism in the NRF2 encoding gene was associated with AMD risk [23]; thirdly, studies have highlighted its potential protective effects on retinal cells. AAV-*Nrf2* showed ability to induce antioxidant gene expression and protected RPE from oxidative injury and photoreceptor dysfunction on a mouse model of oxidative injury with NaIO_3_ intravitreal injection [24] and facilitated structure preservation and retinal damages functional recovery on a light induced mouse model of AMD [25]. However, whether NRF2 directly regulates lysosomal biogenesis or lysosomal reformation remains to be clarified.

Given its importance and potential utility as a therapeutic target, a variety of drugs acting as NRF2 inducers are under development. Dimethyl fumarate (DMF), first approved for the treatment of relapsing-remitting multiple sclerosis, rapidly interacts with cys-151 of the NRF2 regulator, KEAP1, to activate the NRF2 pathway [19]. Some studies have already highlighted DMF protective capacities after light-induced retinal damage [26,27]. In this study, we explore the role of DMF-driven NRF2 activation in antioxidant and autophagy mechanisms in RPE cells, highlighting the value of NRF2 as a therapeutic target of lipofuscin clearance and related retinal degenerative diseases. Our findings demonstrate that macroautophagy plays a key role in the clearance of AFGs and presumably lipofuscin *in vivo*.

Here we have uncovered evidence suggesting a lipofuscin cycle, formation, maintenance and degradation, whereby i) lysosomal failure creates AGFs (a surrogate for lipofuscin), then ii) AFGs loose connection with the endolysosome system and subsequently iii) AFGs can only be degraded by the macroautophagy pathway. NRF2 appears to be one important activator of macroautophagy efficiently clearing AFGs from RPE cells.

## RESULTS

### DMF induces antioxidant and autophagy mechanisms via NRF2

To study the potential role of DMF-driven NRF2 activation in retinal neurodegenerative diseases, we generated a human immortalised RPE cell line (ARPE-19), where the NRF2 encoding gene (*NFE2L2*) was knocked-out (KO) using CRISPR/Cas9-based gene editing (NRF2 KO cells) to compare with wild-type (WT) ARPE-19 (Supplementary Figure S1). Both WT and NRF2 KO cell protein levels were assessed upon incubation with 5 gM DMF (Figure1A). Indeed, 5 gM DMF rapidly induces NRF2, with activation detectable as early as 2h post-treatment, as assessed by Western blot (Figure 1B and Supplementary Figure S2). NRF2 levels peak around 4h and then lower to baseline levels after 8h of incubation. In NRF2 KO cells, no specific NRF2 bands are observed, confirming the KO. DMF-driven NRF2 activation is supported by the induction of HO-1, a canonical NRF2 target, reaching its highest expression upon 8h of incubation (Figure 1A-B and Supplementary Figure S2). DMF appears to induce NRF2-dependent autophagy, as indicated by increased LC3 II/I ratio levels (starting at 2h incubation) in the WT cells, a key autophagy marker (Figure 1A-B and Supplementary Figure S2). Another NRF2 target, p62 (linked to autophagy), shows no significant response to DMF treatment under these conditions. However, increased basal levels of p62 are verified in NRF2 KO cells, possibly because the lack of NRF2 perturbs autophagic flux [28,29] (Figure 1A-B and Supplementary Figure S2).

**Figure 1.**
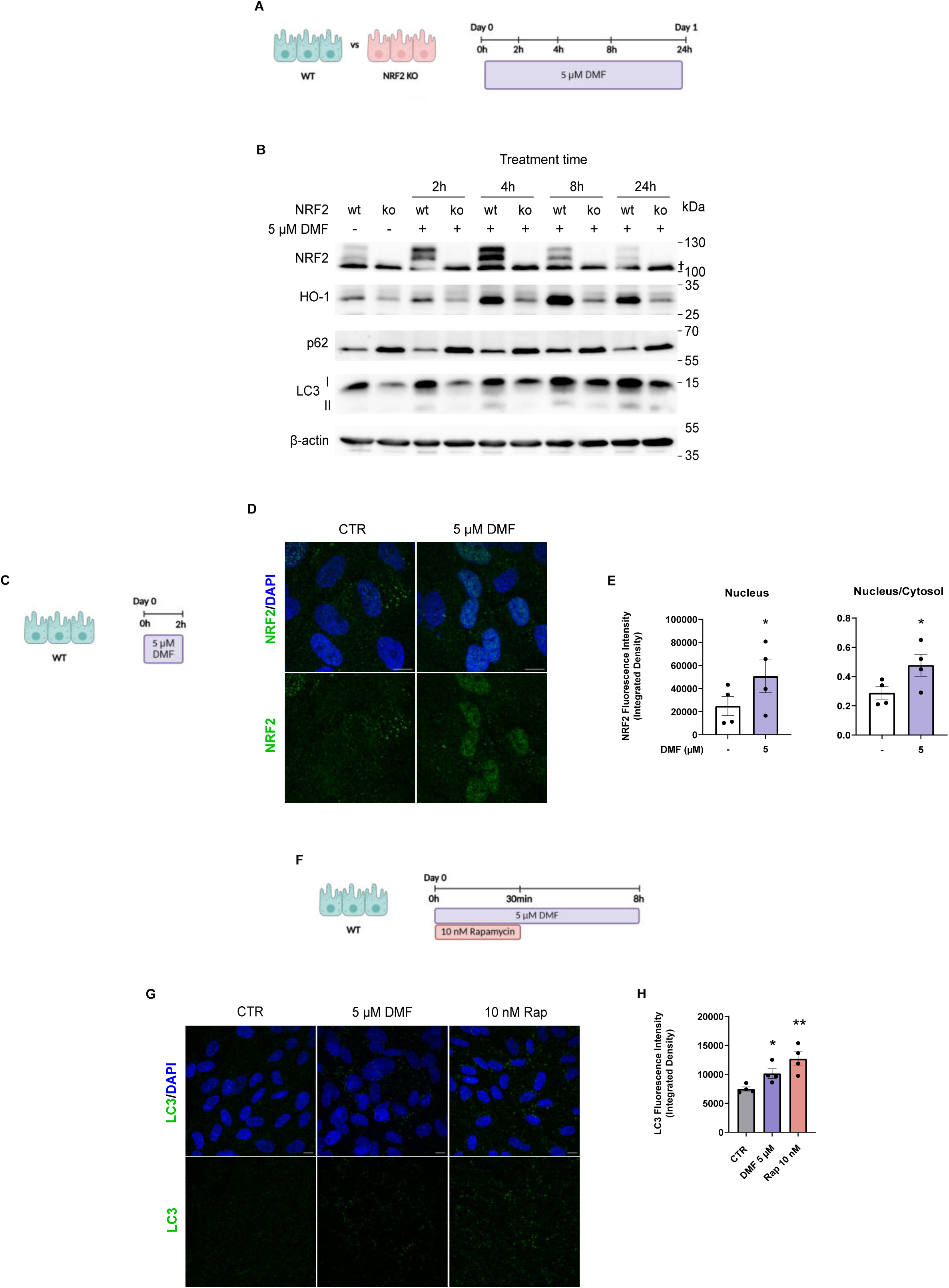
DMF induces antioxidant and autophagy mechanisms via NRF2. (A) WT and NRF2 KO cells treated with 5 gM of dimethyl fumarate (DMF, represented in purple) for 24 hours. (B) Western Blot of NRF2, HO-1, LC3 and p62 protein levels on WT and NRF2 KO cells. † Unspecific band. (C) WT cells treated with 5 gM of DMF (purple) for 2 hours. (D) Immunofluorescence of NRF2 (green) in WT cells treated and non-treated with 5 gM DMF for 2 hours. Nuclei stained with DAPI (blue). Scale bar 10 gm. (E) Quantification of the intensity of Nuclear and the ratio Nucleus/Cytosol NRF2 puncta in WT cells treated and non-treated with 5 gM DMF for 2 hours. Data is presented as Mean ± SEM, representative of n=4 independent experiments. Statistical comparisons were performed using unpaired student’s t-test. * p < 0.05 vs non-treated cells. (F) WT cells treated with 5 gM of DMF (purple) for 8 hours or 10 nM Rapamycin for 30 minutes. (G) Immunofluorescence of LC3 (green) in WT cells treated and non-treated with 5 gM DMF for 8 hours or 10 nM Rapamycin for 30 minutes. Nuclei stained with DAPI (blue). Scale bar 10 gm. (H) Quantification of the intensity of LC3 puncta in WT cells treated and non-treated with 5 gM DMF for 8 hours or 10 nM Rapamycin for 30 minutes. Data is presented as Mean ± SEM, representative of n=4 independent experiments. Statistical comparisons were performed using unpaired student’s t-test. * p < 0.05, ** p < 0.01 vs non-treated cells.

To further support NRF2 and LC3 activation upon DMF treatment, we analysed RPE cells by fluorescence microscopy (Figure 1C-H). After 2h of treatment with 5 gM DMF we observed significantly increased nuclear and nucleus/cytosol ratio levels of NRF2 in WT cells (Figure 1D-E). Additionally, after 8h of treatment with 5 gM DMF we observed a significant accumulation of LC3 signal (Figure 1F) albeit weaker than when RPE cells were treated with 10 nM Rapamycin for 30 min, used as positive control (Figure 1G-H). Rapamycin was previously reported to induce LC3 levels and autophagy [30–32]. These results led us to conclude that DMF treatment in RPE cells was able to enhance both antioxidant and autophagy mechanisms via NRF2.

### NRF2 is activated upon POS phagocytosis

In previous work, we optimised a cellular model mimicking lipofuscin accumulation with AFG formation by challenging RPE cells with a single pulse of purified porcine POS [7]. In the present study, we used this *in vitro* assay to explore if there is a link between AFG formation and NRF2 signalling.

To explore these effects, monolayers of RPE cells were fed with POS for 4h (Figure 2A) and then intracellular ROS levels and NRF2 protein levels were analysed up to 48h later. Upon POS phagocytosis, a significant increase of intracellular ROS is verified when compared to non-treated cells (Figure 2B). ROS production was a previously described consequence of POS phagocytosis processes [1,33]. Corroborating these results, endogenous NRF2 activation is observed immediately, as indicated by increased NRF2 protein levels at the 0h time point, ie, after 4h feeding of POS (Figure 2C and Supplementary Figure S3). This early activation is further supported by the subsequent rise in HO-1 protein levels, a canonical NRF2 target, which peaks at 24h post-phagocytosis. Additionally, p62, shows increased levels both at the initial (0h) and 24h time points (Figure 2C and Supplementary Figure S3), suggesting sustained engagement of NRF2-mediated pathways throughout the early and intermediate phases of POS processing. Notably by 48 hours after phagocytosis, NRF2 and associated protein levels return to values similar to those of untreated cells. The immediate NRF2 activation upon POS ingestion implies that the phagocytic process triggers oxidative or metabolic stress signals, which stimulate antioxidant and cytoprotective responses. The sustained increase in HO-1 and p62 supports that NRF2 not only provides acute stress resistance but also facilitates longer-term cellular adaptation.

**Figure 2.**
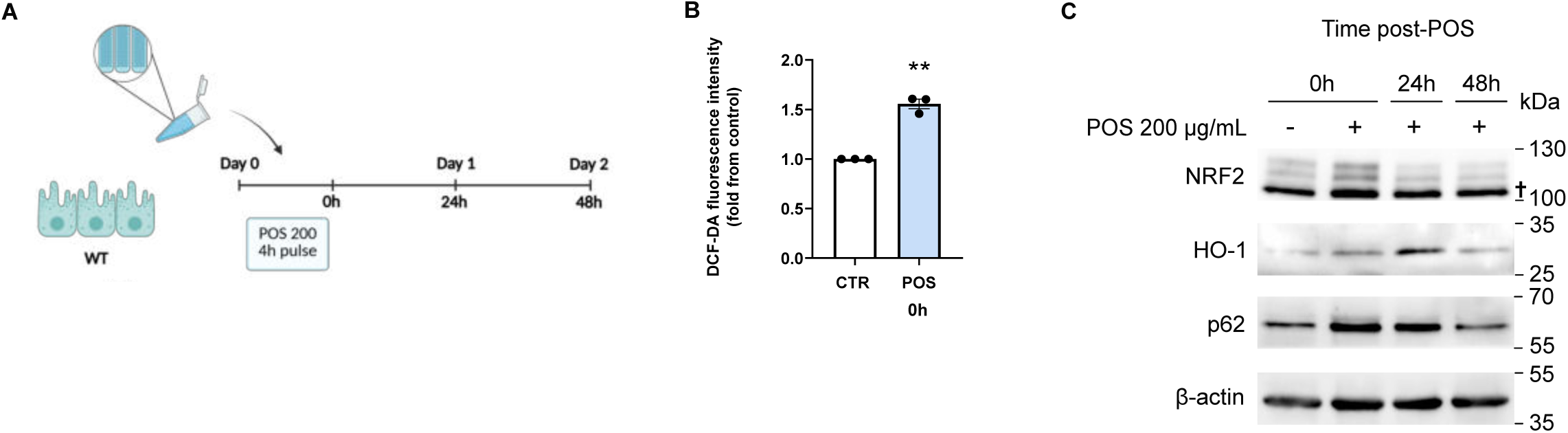
NRF2 is activated upon POS phagocytosis. (A) WT cells incubated 4 hours with POS 200 gg/mL, followed by a chase period of 48 hours (B) Quantification of intracellular ROS levels, immediately after the 4h POS feeding, using the cell permeable and oxidation sensitive DCFDA fluorescent probe. ** p < 0.01 vs No-POS fed cells (control). (C) Western Blot of WT cells incubated 4 hours with POS 200 gg/mL, showing an endogenous activation of NRF2, its target HO-1, and the levels of p62 protein. † Unspecific band.

### AFgs loose connection with the endocytic pathway over time

As shown in previous work, AFGs continue to form over time following POS phagocytosis and derive from incompletely digested POS-containing phagosomes [7]. After 3 days, AFGs are surrounded by a single membrane positive for lysosome markers and rhodopsin present in POS [7]. We showed that lysosome-phagosome fusion was required for AFG formation, such that impairment of lysosomal pH or catalytic activity enhanced AF accumulation. To complement these studies, we measured the number of AFGs over time showing a gradual accumulation over 7 days (Figure 3B and D). We reasoned that this could be due to a “drop-off” from the lysosomal cycle, given their indigestibility. To test this hypothesis, we fed RPE cells with dextran at different times after POS treatment to ask whether the AFGs remain accessible to the endolysosomal pathway over a period of 7 days. As shown in Figure 3B and C, the co-localization of dextran with AFGs decreases significantly over time, suggesting that these AFGs gradually loose connection with the endolysosomal pathway. We suggest they remain in the cytoplasm as “storage granules”.

**Figure 3.**
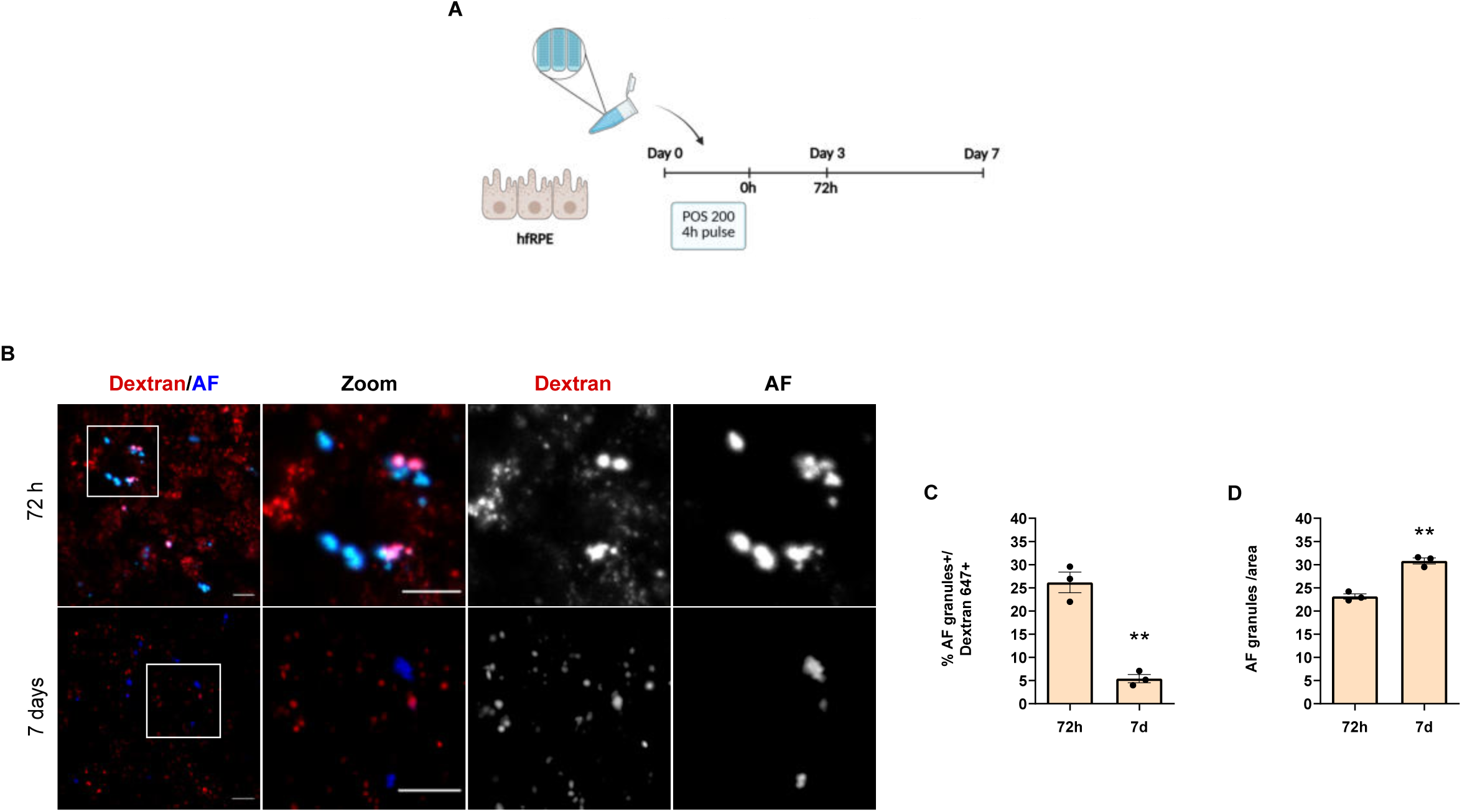
AFgs exit the endocytic pathway over time. (A) hfRPE cells incubated 4 hours with POS 200 gg/mL and chased for a period of 72 hours and 7 days. (B) Live cell imaging of hfRPE cells containing AF (blue) pulsed overnight with Dextran Alexa Fluor 647 (red), washed to remove non-internalized dextran and chased for 4h. Representative images of n=3 independent experiments are shown on the left panel. Scale bar 20 uM, with magnification (zoom), scale bar 10 uM. (C) Quantification of the % of AFG that co-localized with Dextran 647. Data is presented as Mean ± SEM, representative of n=3 independent experiments. Statistical comparisons were performed using paired student’s t-test. ** p < 0.01 vs 72h chase period. (D) Quantification of the number of AFG per area. Data is presented as Mean ± SEM, representative of n=3 independent experiments. Statistical comparisons were performed using paired student’s t-test. ** p < 0.01 vs 72h chase period.

### Pre-treatment and late NRF2 induction with DMF reduce AFg load

Next, we sought to explore the effects of DMF-driven activation of NRF2 on RPE AFG load. Therefore, monolayers of RPE cells were fed with POS for 4h, followed by three different experimental approaches: i) pre-treatment at - 16h; ii) treatment at 0h and iii) treatment at 48h post POS phagocytosis assays (Figure 4A). The quantification of AFG was accessed by flow cytometry (Figure 4B).

**Figure 4.**
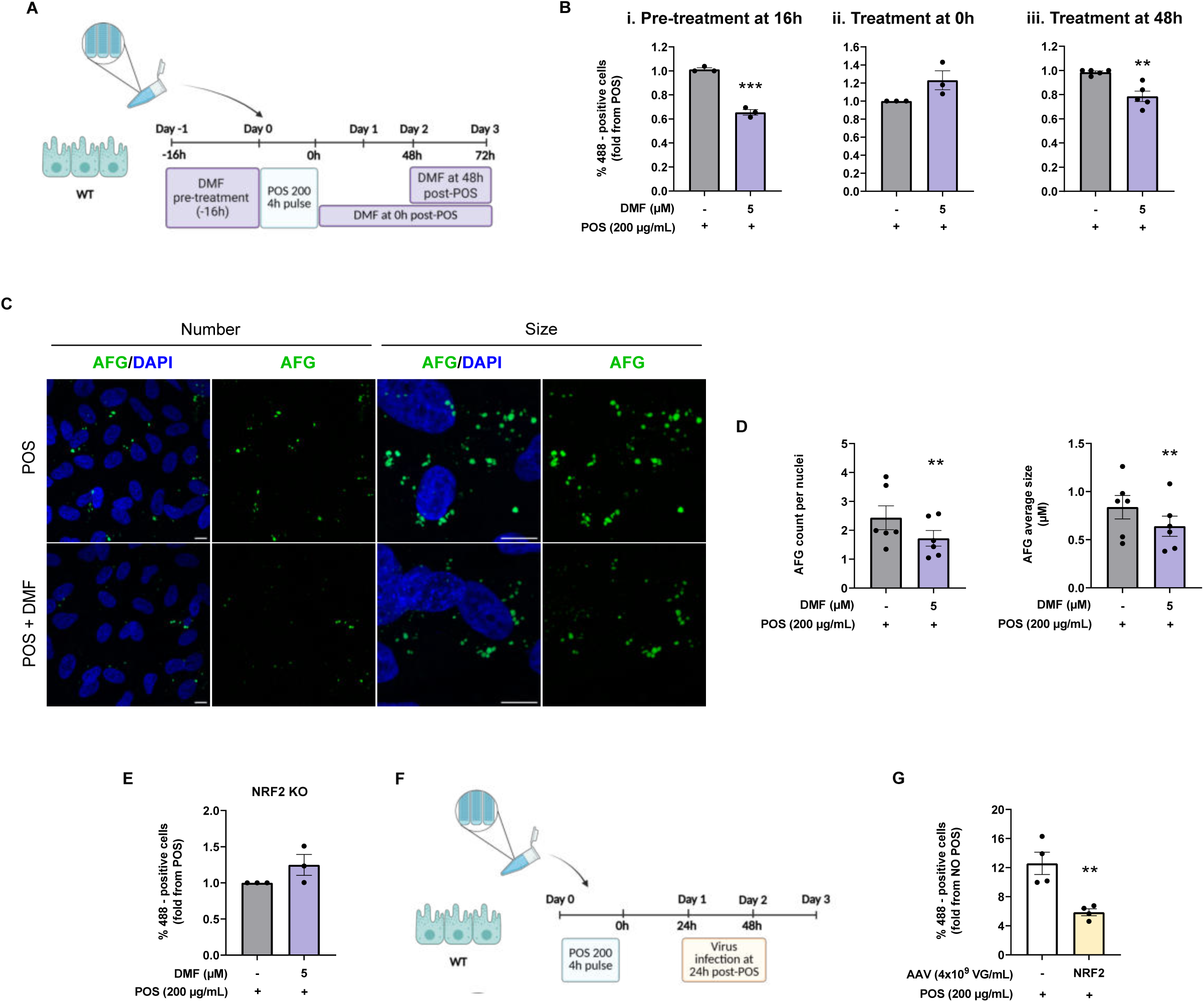
Late NRF2 induction with DMF reduces autofluorescent granules (AFgs) load. (A) WT cells pre-treated 16h with 5 gM of dimethyl fumarate (DMF, in purple) and then incubated 4 hours with POS 200 gg/mL (i) or posteriorly treated with 5 gM of dimethyl fumarate (DMF, in purple) at 0 hours (ii) or at 48 hours (iii) post POS phagocytosis assays. (B) Quantification of % of WT cells containing AFGs by flow cytometry. Data is presented as Mean ± SEM, representative of n=3 independent experiments. Statistical comparisons were performed using unpaired student’s t-test. ** p < 0.01 vs POS fed cells. C) Immunofluorescence of AFGs (green) in WT cells treated and non-treated with 5 gM DMF at 48 hours (ii) post POS phagocytosis assays. Nuclei stained with DAPI (blue). Scale bar 10 gm. D) Quantification of the number of AFGs per nuclei and AFGs size (gm) in ARPE-19 cells treated and non-treated with 5 gM DMF at 48 hours (ii) post POS phagocytosis assays. Data is presented as Mean ± SEM, representative of n=6 independent experiments. Statistical comparisons were performed using paired student’s t-test. ** p < 0.01 vs POS fed cells. (E) Quantification of % of NRF2 KO cells containing AFGs by flow cytometry. Data is presented as Mean ± SEM, representative of n=3 independent experiments. Statistical comparisons were performed using unpaired student’s t-test. No significant differences were seen. (F) WT cells were first pulsed with POS for 4h, then transduced with NRF2-AAVs and followed by a 72h chase period. (G) Quantification of % of WT cells containing AFGs by flow cytometry. Data is presented as Mean ± SEM, representative of n=4 independent experiments. Statistical comparisons were performed using unpaired student’s t-test. ** p < 0.01 vs POS fed cells.

Our results show that pre-treatment at −16h and treatment at 48h after POS phagocytosis with 5 gM of DMF similarly induce a significant reduction in AFGs, while treatment at 0h results in a slight increase in AFG load, assessed by flow cytometry (Figure 4Bii). Autofluorescence intensity following DMF treatment at 48h led to a 25-30% reduction in cellular AFGs (Figure 4Biii). Confocal microscopy imaging and quantification of the number and size of AFGs at 72h post-phagocytosis showed a decrease in number (about 25-30%) and size of the AFGs within RPE cells (Figure 3 C-D).

Several factors may explain these contrasting outcomes: firstly, DMF is a fast-acting compound, described with cellular effects typically evident within 24h (Figure 1B) [7,34]; secondly, POS phagocytosis itself induces endogenous NRF2 activation, as shown in Figure 2C. Therefore, immediate post-phagocytosis DMF treatment (0h) may not work properly as induction of NRF2 is already promoted by phagocytosis of POS; thirdly, the effect observed after DMF treatment at 48h is likely due to inducing a new cycle of DMF activation given the low basal endogenous NRF2 (Figure 2C) and DMF treatment; fourthly, the reduction in AF intensity upon pre-treatment of DMF could be driven by a peak of NRF2 transcription targets being present at the time of phagocytosis, leading to a more efficient clearance of POS-containing phagosomes.

As a control in the phagocytosis assays, we employed NRF2 KO ARPE-19 cells using a 5 gM DMF treatment at 48h, and measure AFGs (Figure 4E). Upon NRF2 depletion, DMF is no longer able to reduce AFG levels (Figure 4E), confirming that these effects are in fact induced via NRF2. Supporting this protective role of NRF2, AAV-mediated overexpression of NRF2 similarly decreases AFG levels (Figure 4F-G), reinforcing the idea that controlled NRF2 activation enhances the degradation and clearance of phagocytosed material.

### AFgs resolution mediated by NRF2 activation depends on autophagy

Finally, we explored the mechanisms of AFG resolution. The experiments above and the literature on NRF2 mechanism of action led us to study the importance of the autophagy pathway on this process. We generated a human immortalised RPE cell line (ARPE-19) where ATG7 was knocked-out using CRISPR/Cas9-based gene editing (ATG7 KO cells) (Supplementary Figure S4). We then performed POS phagocytosis assays as above using a 5 gM DMF treatment at 48h and measure AFGs by flow cytometry (Figure 5A). We observed that the effect of DMF in reducing the AFGs load was completely lost in ATG7 KO cells as compared to WT cells (Figure 5B) and as observed for the NRF2 KO RPE cells (Figure 4E). This finding strongly supports the dependence on functional autophagy in the clearance of AFGs.

**Figure 5.**
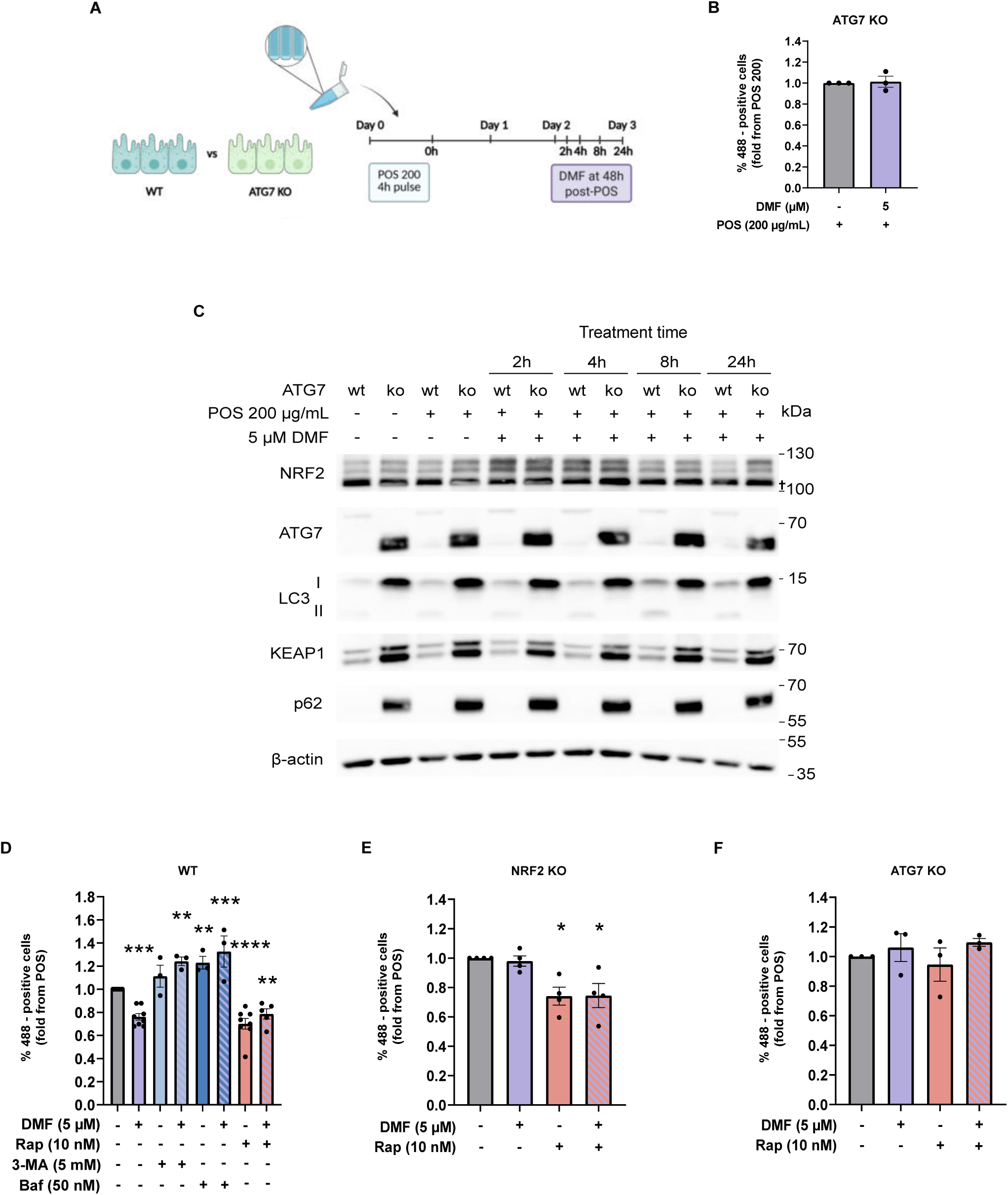
AFgs resolution by mediated NRF2 activation depends on autophagy. (A) WT and ATG7 KO cells incubated 4 hours with POS 200 gg/mL and posteriorly treated with 5 gM of dimethyl fumarate (DMF, in purple) at 48 hours post POS phagocytosis assays. (B) Quantification of % of ATG7 KO cells containing AFGs by flow cytometry. Data is presented as Mean ± SEM, representative of n=3 independent experiments. Statistical comparisons were performed using unpaired student’s t-test. No significant differences were seen. (C) Western Blot of LC3, ATG7, KEAP1 and p62 protein levels on WT and ATG7 KO cells. † Unspecific band. (D-F) Quantification of % of WT (D), NRF2 KO (E) and ATG7 KO (F) cells containing AFGs by flow cytometry, upon exposure to autophagy inducers (Rapamycin, Rap) and inhibitors (3-Methyladenine, 3-MA and Bafilomycin, Baf). Data is presented as Mean ± SEM, representative of n=3 and n=8 independent experiments. Statistical comparisons were performed using one-way ANOVA followed by multiple comparisons Dunnet post hoc correction * p < 0.05, ** p < 0.01 *** p < 0.001, **** p < 0.0001 vs POS fed cells.

Next, we explored the mechanisms of DMF action in the autophagy-deficient cells (Figure 5C and Supplementary Figure S5). ATG7 KO cells exhibit elevated basal expression levels of NRF2, KEAP1, p62, and LC3-I, regardless of DMF treatment. Over time, prolonged NRF2 activation is observed up to 24h, accompanied by sustained high expression of p62 and KEAP1. This pattern is consistent with the non-canonical activation of NRF2, where p62-mediated degradation of KEAP1 enables NRF2 to translocate to the nucleus and drive NRF2 target gene expression [29]. LC3-I is also upregulated in ATG7 KO cells. LC3 accumulates predominantly in its non-lipidated LC3-I form since the lipidation step to generate LC3-II requires ATG7 activity [35,36]. Thus, the detection of increased LC3 −I protein not only confirms the effective knockout but also underlines the disruption of autophagic flux.

Given the clear results using the ATG7 KO line, we wished to confirm the importance of autophagy in the clearance of AFGs. We decided to modulate the autophagic flux and to do so, we used well-tested drugs: one autophagy inducer (Rapamycin) and two autophagy inhibitors (Bafilomycin and 3-MA).

Firstly, we used WT cells (Figure 5D). DMF alone led to a reduction of AFG load as predicted (Figure 4). Following treatment with rapamycin, we observed a reduction in AFG load whether or not DMF was also present. Indeed, the effect of rapamycin and the mTOR/TFEB pathway in reducing the load of AFGs was previously explored using the same *in vitro* assays [16]. Conversely, treatment with autophagy inhibitors led to the loss of this effect. In fact, the AFG load increased whether or not DMF was present. This effect is presumably due to an extensive lysosomal failure blocking degradation (Figure 5D).

Secondly, we used NRF2 KO cells (Figure 5E). As expected, DMF loses its ability to reduce the AFG load (Figure 4). The autophagy activator rapamycin had the effect of reducing the AFG load, as expected [16], in a NRF2 independent way (Figure 5E).

The data shown suggests that the mTOR/TFEB pathway and the NRF2 pathway are independent but use macroautophagy as a common mechanism for the final degradation of AFGs. To test more directly this hypothesis, we used ATG7 KO cells (Figure 5F). The results clearly show that neither DMF nor rapamycin are able to reduce the load of AFGs in ATG7 KO cells. We postulate that ATG7 activity, presumably through macroautophagy is required for both NRF2 and mTOR/TFEB pathways to be effective in clearing AFGs from cells.

### Viral NRF2 over-expression protects RPE cells *in vivo*

The most common disease of the eye in the western world is AMD [1]. In AMD, the key pathogenic mechanism appears to be the degeneration of the RPE caused by multiple factors, among which are oxidative stress, lipofuscin, drusens, the complement pathway and inflammation. In this context, the NRF2 pathway could be beneficial as it is well known to activate anti-oxidative stress responses, and we show here that it is effective in the removal of toxic lipofuscin.

As a proof of concept of the protective effective of NRF2 *in vivo*, we utilized a commonly used model of RPE degeneration and to a certain extent AMD, namely the intra-peritoneal injection of the powerful oxidant NaIO_3_ [37], after sub-retinal injection of AAV-NRF2 (treated eye) (Supplementary Figure S6). As a proper control to the treated eye, we used a saline solution by sub-retinal delivery in the contralateral eye (Figure 6, upper panels). We observed that one single intraperitoneal injection of NaIO_3_ is able to induce loss of RPE cytoarchitecture. This loss of RPE is observed after 5 days post-injection and is more notable in the centre of the eye (although mice have no macula). The transition zone is full of altered RPE cells with fusiform and other shapes and no longer forming an epithelium (Figure 6, upper panels). The peripheral areas are commonly protected and the structure maintained with the relatively mild concentration of NaIO_3_ we use. To test the potential protective effect of the NRF2 pathway including the ability to clear lipofuscin, we injected via sub-retinal delivery an AAV-driven vector over-expressing NRF2 and analysed RPE cells as above (Figure 6, lower panels). We observed that the over-expression of NRF2 in the RPE effectively prevented cell damage in the transduced areas following NaIO_3_ administration. We conclude that pharmacological activators of NRF2 may be beneficial counteracting the age and disease-driven degeneration of the RPE.

**Figure 6.**
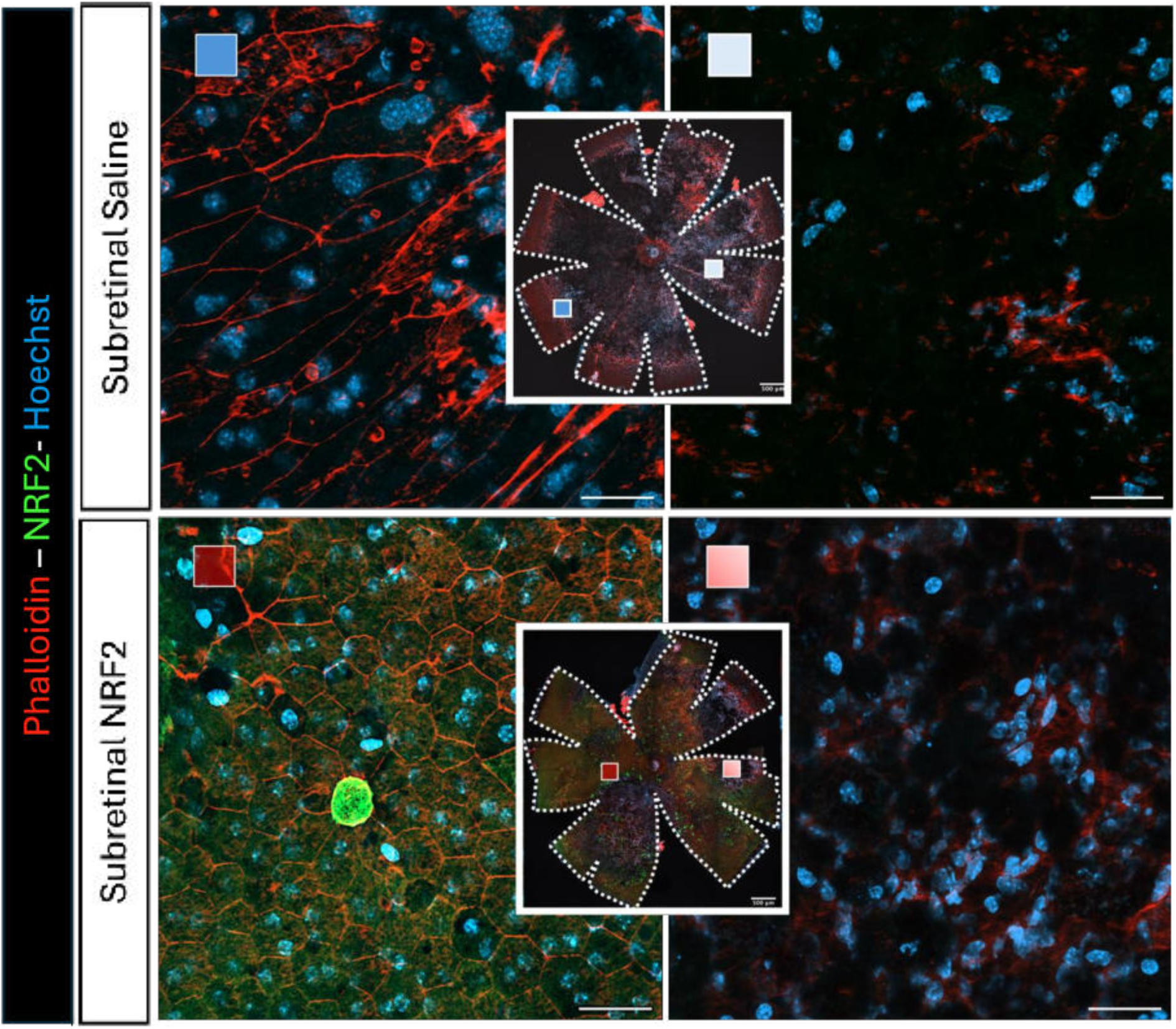
NRF2 overexpression preserves RPE cytoarchitecture in an induced mice model of RPE degeneration. Control Injected Eye (upper panels). Subretinal injection of saline followed by NaIO_3_ injection exhibits extensive RPE loss and architectural disruption, as indicated by diminished and disorganised Phalloidin (red) staining, confirming NaIO₃-induced RPE degeneration (central square, scale bar 500 μm). Magnification shows the area of elongated and hypertrophic RPE cells in the peripheral zone (dark blue square). A magnification reveals pronounced RPE cell loss and structural breakdown (light blue square). Scale bar 30 μm. AAV-NRF2-treated eye (lower panels). Subretinal injection of AAV-NRF2 followed by NaIO_3_ shows preservation of the RPE monolayer, with intact Phalloidin (red) staining exclusively in regions exhibiting NRF2 (green) expression (central square, scale bar 500 μm). A magnification image from the equatorial region (dark red square) demonstrates a strong spatial correlation between AAV-NRF2 expression and maintenance of normal RPE morphology. A magnification view of an untransduced region lacking the NRF2 signal shows loss of RPE integrity, similar to the control NaIO_3_ injected eye (light red square). Scale bar 30 μm.

## Discussion

Autophagy impairment and lysosomal dysfunction in RPE cells are central pathological mechanisms of retinal neurodegenerative diseases [1,7]. Here we show that a critical mechanism for the clearance of AFGs (and presumably lipofuscin *in vivo*) is macroautophagy. Furthermore, we show that the NRF2 and mTOR/TFEB pathway activation act in the clearance of AFGs as inducers of macroautophagy. Therefore, modulation of macroautophagy by these two pathways or by other means opens a potentially new therapeutic strategy to help preserve RPE health in early and intermediate AMD.

In this study, we focus on the activation of the NRF2 pathway in RPE cells challenged with large amounts of POS and study its role in the formation and degradation of resulting AFG granules mimicking lipofuscin accumulation *in vivo*. We first uncover a novel feedback mechanism, whereby POS phagocytosis endogenously triggers NRF2 activation, suggesting a natural link between cellular debris and phagocytic activity to the activation of protective stress responses (Figure 2). A similar effect was observed for the mTOR/TFEB pathway in the same *in vitro* model [16]. This physiological NRF2 induction most likely serves as an adaptive mechanism to neutralize oxidative stress associated with daily photoreceptor turnover, trying to protect RPE cells from oxidative damage and maintaining cellular function under normal conditions [1,38]. Thus, NRF2 is an important oxidative stress sensor during POS processing, a role that remains largely unexplored.

The FDA-approved drug DMF activates NRF2 by allowing newly synthesised NRF2 to translocate into the nucleus and initiate transcription of target genes [39]. We first showed that DMF activates the NRF2/HO-1 pathway in RPE. The antioxidant effects of DMF in retinal cells have been recently confirmed in human retinal endothelial and ganglion cells, showing cytoprotection against oxidative stress and promotion of cell survival via NRF2/HO-1 pathway activation [40,41]. These findings indicate that pharmacological NRF2 activation by DMF can enhance intrinsic antioxidant defences in the retina. Furthermore, we verify that these protective effects are extended to the autophagy/lysosomal pathway through increased LC3-II (Figure 1). Increased LC3 protein as a result of DMF treatment was also recently observed in murine microglia cells [28,42]. This interpretation is also supported by the reported increase of p62 upon NRF2 depletion (Figure 1). Given that a feedback loop has already been established between NRF2 and p62 [29], the loss of NRF2 likely disrupts this regulatory balance, which may lead to dysregulated or compensatory changes in p62 levels, thereby underlying the importance of the NRF2 pathway in maintaining autophagy homeostasis. Similar evidence was published in a different system whereby Nrf2 is required for proper induction of autophagy and that its loss results in a dysfunctional Keap1–Nrf2–p62 feedback loop leading to p62 protein accumulation due to impaired autophagic flux [28].

Lipofuscin is a by-product of incomplete POS degradation, leading to chronic oxidative stress in a self-perpetuating feedback loop [1]. Late or pre-injury NRF2 induction by DMF results in decreased AFG load, a hallmark of impaired lysosomal degradation, underlining the therapeutic potential of transient NRF2 activation and indicates improved autophagic clearance capacity in RPE cells. Pharmacological enhancement of NRF2 signalling maintains antioxidant defences and autophagy over time, which can reverse or avoid lipofuscin formation and accumulation associated with retinal aging and degenerative diseases. This supports the concept of NRF2 not just as an acute stress responder but as a regulator of long-term retinal health. Consistently, transient NRF2 activation triggers endogenous defence programs that ultimately strengthen the organism’s resilience and promote longevity [21,43]. Autophagy facilitates degradation of photoreceptor components and prevents accumulation of lipofuscin-like aggregates [31,44]. As we show here, the beneficial effects of DMF are dependent on functional autophagy pathways, as both genetic and pharmacological ablation of autophagy abolish these improvements. Our results indicate that ATG7-dependent autophagy is crucial for effective clearance of AFGs and demonstrate that the interplay between NRF2 and autophagy is a critical component of AFG resolution (Figure 5). NRF2-mediated AFG clearance depended on functional autophagy emphasizes that antioxidant signalling alone is insufficient; instead, NRF2 activation supports autophagic flux, possibly via transcriptional upregulation of autophagy-related genes and maintenance of the redox environment necessary for lysosomal enzyme function [1,45]. This crosstalk promotes cellular clearance and prevents accumulation of potentially cytotoxic waste products.

Importantly, the therapeutic potential of NRF2 gene delivery in retinal degeneration has been further reinforced by the recent study of Gardner and colleagues [46], who demonstrated that AAV-mediated NRF2 gene therapy preserves retinal structure and function in rodent models of oxidative damage. The authors demonstrated significant protection of both the RPE and photoreceptors following NaIO_3_-induced oxidative injury in rats and mice. Notably, this protection extended beyond morphological preservation, with maintenance of retinal architecture assessed by optical coherence tomography (OCT) and preservation of visual function assessed by electroretinography (ERG). These findings provide compelling independent evidence supporting NRF2 gene delivery as a promising therapeutic strategy against oxidative stress-associated retinal degeneration. Our findings complement and extend these observations by providing a potential cellular mechanism underlying the protective effects of NRF2 gene therapy. While Gardner et al. primarily demonstrated the efficacy of sustained AAV-mediated NRF2 expression in preserving retinal structure and function, our study identifies an additional dimension of NRF2-mediated protection: the regulation of autophagy-dependent clearance of lipofuscin-like AFGs in RPE cells. Specifically, we demonstrate that NRF2 activation promotes AFG resolution through an ATG7-dependent mechanism, suggesting that NRF2-mediated retinal protection may involve not only the enhancement of antioxidant defences but also the improvement of cellular proteostasis and clearance of potentially toxic intracellular material. These effects are particularly relevant in the context of AMD, where chronic oxidative stress and progressive accumulation of undegraded material contribute to RPE dysfunction.

The finding that NRF2 is physiologically activated during POS phagocytosis can be further enhanced pharmacologically by DMF and the cell protection effect of over-expressing NRF2 *in vivo* present a compelling rationale for repurposing DMF and/or other activators of NRF2 as an AMD therapeutic, especially considering its established safety profile in other chronic diseases. Our *in vitro* and *in vivo* results lay the groundwork for future translational and *in vivo* studies assessing the efficacy of DMF or similar NRF2 activators in preventing or slowing the progression of retinal degeneration.

In summary, this work presents fundamental and translational novelties (Figure 7). On a fundamental side (Figure 7A), it reveals the full life cycle of lipofuscin-like AFGs. POS phagocytosis can be overwhelming to RPE cells, causing lysosomal dysfunction and creation of AFG organelles. These AFGs mature and loose connection with the endolysosomal pathway becoming “storage” granules causing chronic oxidative stress in the RPE presumably through lipoperoxidation. This process is particularly relevant to retinal ageing, as the post-mitotic nature of RPE cells limits the clearance of accumulated intracellular material through cell division, contributing to the progressive accumulation of lipofuscin with age. Upon macroautophagy activation by several forms including the activation of NRF2 and mTOR/TFEB pathways, a significant portion of AFGs can be cleared in autolysosomes (Figure 7A). Many details are missing which should be a focus of future research. On a pre-clinical side (Figure 7B), the overall findings that NRF2 is physiologically activated during POS phagocytosis, can be further enhanced pharmacologically by DMF and the cell protective effect of over-expressing NRF2 *in vivo*, present a compelling rationale for repurposing DMF as a retinal therapeutic (Figure 7B), especially considering its established safety profile in other chronic diseases. Future studies should address long-term efficacy and safety of DMF (and other NRF2 activators) in preclinical and clinical settings. Ultimately, this work advances the mechanistic understanding of retinal pathogenesis and underscores the therapeutic potential for innovative activators of macroautophagy such as NRF2 activators in early and intermediate AMD as well as other chronic age-related diseases.

**Figure 7.**
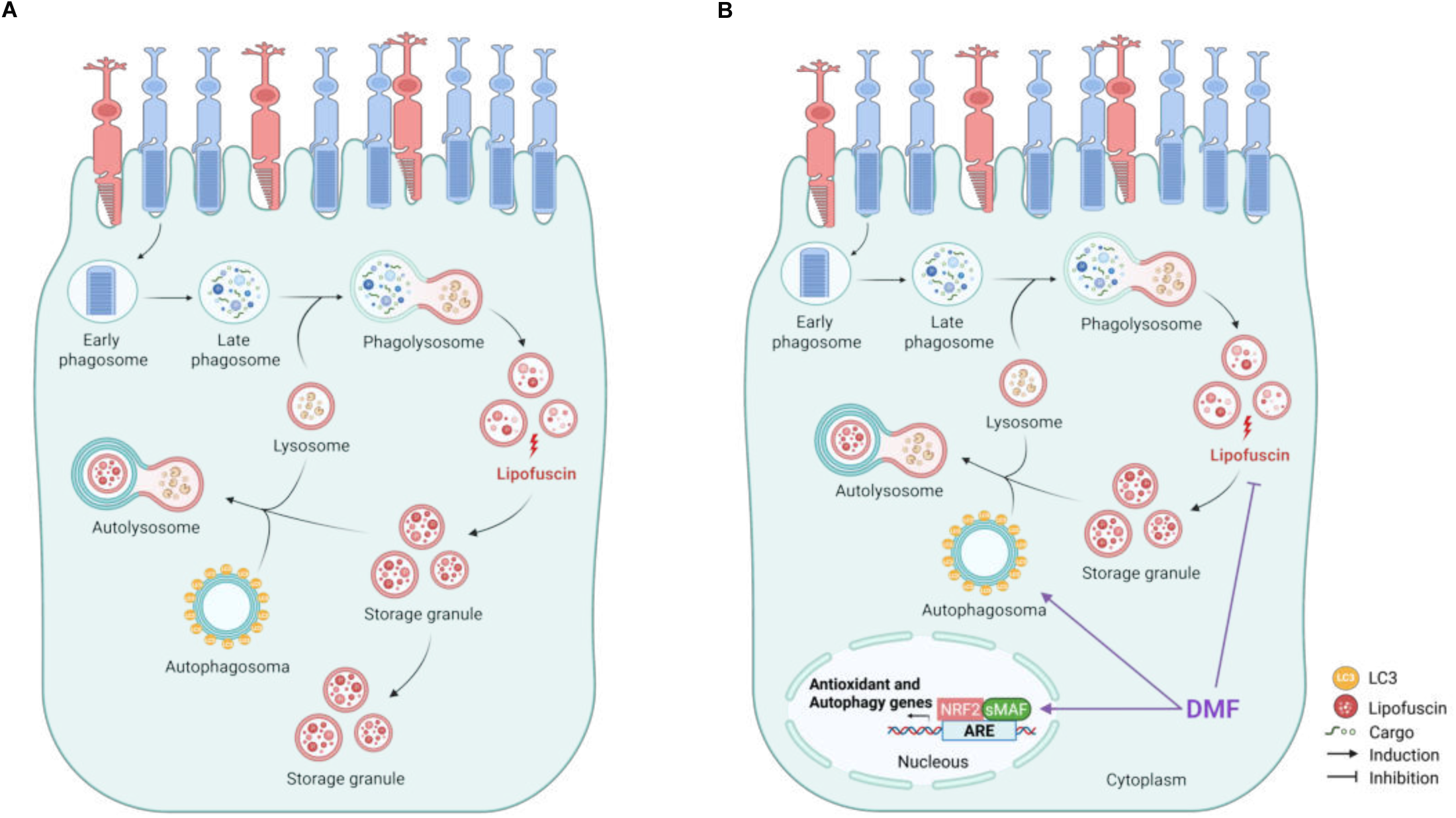
Proposed model of DMF–NRF2-driven autophagic clearance of AFgs in RPE cells. (A) Retinal pigment epithelium (RPE) cells phagocytose shed photoreceptor outer segments (POS), which are processed through the phagosome-lysosome pathway. Under conditions of lysosomal dysfunction, incomplete lysosomal degradation of this material generates lipofuscin, an undegradable by-product that accumulates in storage granules, that could be not fully cleared via macroautophagy, (B) DMF treatment rapidly activates NRF2 signalling: NRF2 translocates to the nucleus, heterodimerizes with sMAF, and binds the antioxidant response element (ARE) to induce transcription of antioxidant and autophagy-related genes. This upregulates autophagosome formation and autophagic flux, increasing sequestration and degradation of lipofuscin-containing storage granules through the autophagosome–lysosome fusion, thereby enhancing clearance of accumulated lipofuscin via macroautophagy.

## Methods

### Cell Culture

ARPE-19 cells (ATCC, Manassas, VA) were cultured in Dulbecco’s Modified Eagle Medium/Nutrient Mixture F-12 (DMEM/F-12) (Gibco, Grand Isle, NY) supplemented with 10 % fetal bovine serum (FBS) (Gibco) and 1 % penicillin–streptomycin (Pen/Strep) (Gibco). Primary human fetal RPE cells (hfRPE) were purchased from Lonza and cultured using optimized RtEBM^TM^ Basal Medium and RtEGM^TM^ SingleQuots^TM^ Supplements (Lonza), as already described [7].

### ARPE-19 gene editing

ARPE-19 NRF2 and ATG7 knock-out (KO) cell lines were generated with genome editing. TrueDesign™ Genome Editor online tool from ThermoFisher Scientific ™ was used to design the single guide RNA (sgRNA). The selected sgRNAs (5’ GATATTCCCGGTCACATCGA 3’ and 5’ GAAGCTGAACGAGTATCGGC 3’) targeted the NFE2L2 exon 4 and the ATG7 exon 1, respectively. The KOs were generated using the Alt-R CRISPR-Cas9 system (Integrated DNA Technologies, IDT) following electroporation with the Neon Transfection System Kit (#MPK1025K, 10 gL, Invitrogen) protocol. Briefly, low passage, ARPE-19 cells were grown to 70 % confluence in a 25 cm^2^ t-flask. When confluency was reached the cell media was changed. The next day, 5.6 x 10^5^ cells were resuspended in 9 gL of Resuspension Buffer (Buffer R, Neon Transfection). The designed sgRNAs, HS. Cas9. NFE2L2.1. AB (#237499234, IDT) and HS. Cas9. ATG7.1. AA (#229767851, IDT) were diluted to a final concentration of 44 gM. Then, the Ribonucleoprotein (RNP) complex was formed by diluting the Alt-R S.p. Hifi Cas9 Nuclease V3 (#1081060, IDT) in Buffer R for a final concentration of 36 gM, to then mix to the diluted Alt Cas9 enzyme. The complexation reaction was left to progress at room temperature (RT) for 15 minutes. Meanwhile, the 100 gM Alt-R Cas9 Electroporation enhancer (#1075916, IDT) was diluted in molecular water to reach a final concentration of 10.8 gM (working solution). Finally, the RNP complex was added to the cell suspension together with the Alt-R enhancer working solution. Electroporation was then carried out with an intensity of 1 350 mV, along 2 pulses of 20 ms, accordingly to manufacturer’s instructions. Cells were seeded in a 12-well plate with pre-warmed supplemented DMEM with 10 % FBS. When confluency was reached, cells were split onto super-low confluency (1:25, 1:50, 1:100, 1:200, 1:500 and 1:1000). As soon as cells started to form aggregates, different clones were isolated from the different dilution wells into a 48-well plate. Colonies were allowed to expand, and positive clones were confirmed by Western blot and Sanger sequencing.

### POS phagocytosis assays

Before phagocytosis assays, POS isolated in the lab [7] were thawed in the dark, washed three times in PBS by sequential centrifugations at 5 000 rpm for 10 minutes. To mimic lipofuscin accumulation, RPE cells were fed for 4 hours with POS at a final concentration of 200 μg/mL (∼2.6×10^5^ POS particles/cm^2^ or ∼1×10^6^ POS particles/mL) in supplemented DMEM with 10 % FBS and 1% Pen/Strep. After this pulse, POS were removed, cells were washed and cultured in supplemented DMEM with 10 % FBS (or in supplemented DMEM with 1% FBS for hfRPE) and 1% Pen/Strep. To study the mechanisms of POS clearance, the beginning of feeding was considered as 0 h. Cells were later transduced with AAV, treated with DMF or trypsinized for WB analysis during the chase (0, 24 and 48h) periods.For live cell imaging, cells were used after 72h and 7days chase periods.

### DMF treatments

Different experimental approaches were tested: a) For treatment, cells were treated with 5 gM DMF (#242926, Sigma Aldrich) for 2, 4, 8 and 24 hours. Alternatively, after POS phagocytosis assays, cells were treated once with DMF at 0h or 48h after POS feeding; b) For pre-treatment, cells were treated with 5 gM DMF at −16h prior the POS phagocytosis assays. After the incubation periods, the cells were lysed for WB assays, fixed for microscope imaging or trypsinized for flow cytometry assays.

### AAVs transduction

24h after POS phagocytosis assays, ARPE-19 were transduced with Adeno-associated virus serotype 6 (AAV-6) to overexpress NRF2 (AAV-NRF2, construct provided by Boehringer Ingelheim). Cells were incubated with individual AAVs at a MOI of 1×10^9^ VG/well (48-well plate), in OptiMEM at 37°C. After the initial 4h incubation, each well was supplemented with an equal volume of DMEM with 10 % FBS and 1% Pen/Strep. After an additional 20h, AAV-containing media were replaced with fresh supplemented DMEM with 10 % FBS and 1% Pen/Strep. Effects on AF was confirmed by flow cytometry at 72h post-POS phagocytosis assays. Results were represented as the percentage of 488-positive cells normalized to the values of fluorescence detected in cells without the POS pulse.

### Autophagy flux assays

To study autophagic flux, 48 h after POS phagocytosis assays, cells were separately treated for 24h with an autophagy inducer – Rapamycin (Rap, 10 nM, #R8781) and two inhibitors - Bafilomycin A1 (Baf, 50 nM, # B1793) and 3-Methyladenine (3-MA, 5 mM, #M9281), all from SigmaAldrich. Bafilomycin is a late-stage autophagy inhibitor, that functions by selectively blocking vacuolar-type H+-ATPases (V-ATPases) [47] and 3-MA blocks autophagosome formation by inhibiting class III phosphatidylinositol 3-kinases (PI3K) [48]. In parallel, on top of these treatments, cells were also treated with 5 gM DMF to assess the effects of DMF on the autophagic flux. After the incubation period was completed, effects on AF were confirmed by flow cytometry.

### Flow cytometry

RPE cells were trypsinized with TrypLE Express Enzyme (Gibco) for 5 minutes at 37 °C and 5 % CO_2_. Cells were resuspended in supplemented DMEM with 10 % FBS and 1% Pen/Strep, washed with PBS and resuspended in flow cytometry buffer (1 % FBS and 2 mM EDTA in PBS). Data acquisition was performed in a FACS CANTO II flow cytometer (BD Biosiences), with at least 20 000/30 000 cells per condition using the 488 nm excitation wavelength to evaluate cellular AF. Data analysis was performed using FlowJo (Version 10, BD Biosciences). Results were represented as the percentage of 488-positive cells removing the AF background from cells and normalized to the values of fluorescence detected in cells pulsed with POS but in the absence of compound treatment.

### Western blotting

RPE cells were lysed with ice-cold cell lysis buffer (#9803, Cell Signalling Technology) containing Halt Protease and Phosphatase Inhibitor Cocktail (#78444, ThermoFisher Scientific) according to the manufacturer’s instructions. Lysates were pelleted for 15 minutes at 13 000 rpm at 4 °C and protein concentrations were determined with BCA Protein Assay Kit (#23227, ThermoFisher Scientific) in accordance with the manufacturer’s protocol. 30 mg of protein lysate, previously desaturated at 95°C for 5 minutes, were resolved on 15 % and/or 8 % sodium dodecyl sulfate–poliacrylamide gels and subsequently transferred onto 0.2 mm nitrocellulose membranes using Bio-rad Trans-Blot Turbo Transfer System. Membranes were blocked using 5 % of Bovine Serum Albumin (BSA, NZYtech, MB04603) in Tris-buffered saline (TBS, 40 mM Tris, 275 mM NaCl, pH 7.6) containing 0.1 % Tween-20 (SigmaAldrich, 9005-64-5) (TBS-T) before primary antibody incubation overnight, at 4 °C with shaking. The following primary antibodies were used at the indicated dilutions: ATG7 (#376212, Santa Cruz, 1:1 000), GADD34 (#10449-1, Proteintech, 1:1 000), HO-1 (homemade by Cuadradós lab, 1:2 000), KEAP1 (#8047, Cell signaling, 1:1 000), Lamp1 (#3243, Cell signaling, 1:1 000), LC3 (#2775, Cell Signaling,1:1 000), NRF2 (#62352, Abcam, 1:1 000), S6 (#2217, Cell Signaling, 1:1 000), p62 (#28359, Santa Cuz, 1:1 000) and pS6 (#4858, Cell Signaling, 1:1 000). After three washes with TBS-T, HRP conjugated secondary antibodies Sheep anti-Rabbit (#NA934, GE Healthcare, 1:5 000) or Donkey anti-Mouse (#NA931, GE Healthcare, 1:5 000) was incubated for 1 hour at RT. For the case of the anti-β-actin-Peroxidase antibody (#A3854, SigmaAldrich, 1:25 000), membranes were incubated with the antibody for 40 minutes at RT. Protein expression was detected by chemiluminescence using Amersham ECL Prime kit (#RPN2232, Cytiva) or SuperSignal West Atto Ultimate Sensitivity Chemiluminescent Substrate (#A38555, ThermoFisher Scientific) in a Bio-rad ChemiDoc Imaging System.

### Confocal immunofluorescence microscopy

ARPE-19 cells coated in glass coverslips were immediately fixed with 4 % paraformaldehyde (PFA, #43388, Alpha Aesar) for 15 minutes at RT or 100 % methanol for 5 minutes at −20 °C. After fixation, cells were blocked and permeabilized in PBS containing 1 % BSA (#MB04603, NZYtech) and 0.05 % saponin (#47036, Sigma-Aldrich) for 30 minutes. Cells were then incubated with the primary antibody against NRF2 (#12721, Cell Signaling, 1:200) or LC3 (#2775, Cell Signaling, 1:100) for 1 hour, followed by the secondary antibody Alexa Fluor 488 Goat anti-Rabbit (#A11034, Invitrogen, 1:500) for 40 minutes in limited light exposure. Alternatively, cells were incubated with Phalloidin 647 (#A22287, ThermoFisher, 1:400) for 30 minutes. Lastly, cell nuclei were labelled with DAPI (Sigma, 1 μg/mL) and mounted in Mowiol mounting media (Calbiochem). Images were captured in a LSM980 (Zeiss) Confocal Microscope with a Plan-Apochromat 63×1.4 NA oil-immersion objective. Images of NRF2 were obtained with 2.5x zoom. Images of LC3 and AFGs number were obtained with 1.0x zoom. Meanwhile, AFGs size obtained with 3.0x zoom. All images were acquired as Z stacks and analysed using ImageJ software (https://imagej.nih.gov/ij/). Quantifications were carried out using macros or automated segmentation in the ImageJ software.

### Live cell imaging

Three or seven days after POS-phagocytosis assays, hfRPE cells cells plated in 8-well Lab-Tek chamber slides were pulsed with 0.3 mg/mL of Dextran Alexa Fluor 647 (#D22914, Invitrogen), in cell culture medium overnight. Non-internalized dextran was removed by washing the cells three times with PBS. The cells were chased in dextran-free medium for 4h before imaging. Image acquisition was performed using a Zeiss LSM980 confocal microscope, equipped with a 63x 1.4 NA oil objective with 1.5x zoom. All images were acquired as Z stacks and analysed using ImageJ software (https://imagej.nih.gov/ij/).

### ROS measurement

Intracellular ROS levels were assessed using the cell permeable and oxidation sensitive fluorescent probe 5-(and-6)-chloromethyl-2’,7’-dichlorodihydrofluorescein diacetate, acetyl ester (CM-H₂DCFDA, C6827, ThermoFisher Scientific), according to the manufacturer’s protocol. Briefly, 45 minutes prior POS phagocytosis assays, ARPE-19 cells were incubated with 20 gM of freshly prepared CM-H₂DCFDA in phenol red-free DMEM/F-12 medium (Gibco) at 37 °C, with limited light exposure. After incubation, excess of probe was removed by washing with PBS and phagocytosis essays were performed using phenol red-free DMEM/F-12 medium. Upon completion of phagocytosis assays, fluorescence was measured using a Tecan Infinite F200 PRO plate reader with excitation wavelength at 485 nm and emission wavelength at 535 nm. Negative (cells without the probe) and Positive (100, 250 and 500 gM concentrations of hydrogen peroxidase) controls were used. Results were represented as fluorescence intensity removing the fluorescence background from cells without the probe and normalizing to the values of fluorescence detected in cells without the POS pulse.

### Animal experimentation

#### Animals

In this study, all female C57BL/6J mice were used as wild-type (WT) for AAV-NRF2 experiments. All mice were generally used for experiments at 8-10 weeks of age. Mouse colonies were maintained under specific pathogen free conditions in 12/12-hour light/dark cycles, at 21 °C and 50% humidity, at the Champalimaud Research rodent facility. All procedures were conducted in accordance with the Association for Research in Vision and Ophthalmology (ARVO) guidelines and in accordance with the approved national and local institutional committees, namely Direcção Geral de Veterinária (DGAV) and the Champalimaud Animal Care Committee (ORBEA).

#### Experimental groups

All female mice were randomly divided into two groups: Group-1 (Control) did not receive subretinal injections at Day 0 and at Day 1 these mice received intraperitoneal (IP) injections of saline; Group-2 received subretinal injections at Day 0 in the left eyes (AAV-NRF2) and right eyes (saline), using a microinjector. At Day 1 these mice received IP injections of NaIO_3_ (25mg/Kg). Five days after subretinal injection, all eyes were scanned by Optical coherence tomography (OCT) before sacrificing the mice. After dissection of the eyes, flat mounts were immunostained and images were acquired by confocal microscopy.

#### AAV Subretinal Injection

Adult mice (8-10 weeks old) were anaesthetised by intraperitoneal injection of ketamine-xylazine mix (100 mg/Kg ketamine and 10 mg/Kg xylazine), and pupils were dilated with 1 % of tropicamide ophthalmic solution (TropicilTop, 10mg/mL). Subretinal injections were performed under the stereoscope using an RPE-intraocular injection kit connected to an automatic UltraMicroPump3 (UMP3T-2, WPI), delivering AAV-NRF2 suspension at 100 nL/s. A small incision was performed with a 30-gauge needle to puncture a hole slightly below the limbus. Subretinal injections were performed through this incision using a 35-gauge blunt needle coupled to the NanofilTM RPE intraocular injection kit (WPI) to deliver 1.5 μL of AAV-NRF2 into the subretinal space between the retina and RPE, at a concentration of 3.18 × 10‘² vector genomes (VG)/mL. After injections, the eyes were covered with 1 % Carmellose sodium (10 mg/mL, Celluvisc, Allergan) to prevent corneal dehydration during recovery. Optical coherence tomography using Spectralis HRA+OCT (Heidelberg Engineering, Heidelberg) was performed 5 days after the procedure to confirm successful subretinal injections.

#### NaIO_3_ model

To induce retinal oxidative stress, a well-established NaIO₃ model targeting retinal pigment epithelium (RPE) cells was employed [49]. A sterile 2.5 mg/mL NaIO₃ stock solution was freshly prepared by dissolving solid NaIO₃ in 10 mL of sterile 0.9% saline. Group 2 received intraperitoneal (IP) injections of NaIO₃ at a dose of 25 mg/kg one day after subretinal injection (day-1) to induce oxidative stress in the RPE. The control group (Group 1) received an IP injection of an equivalent volume of sterile 0.9% sodium chloride solution on the same day.

#### Confocal immunofluorescence microscopy

Mice were euthanized after 4-5days of subretinal injections by cervical dislocation, and the eyes were enucleated. Eyes are left in 4% PFA for 30 minutes at room temperature after making a puncture in the cornea with a 30G needle. Then, they were washed with PBS containing 0.1% Sodium Azide and transferred to a Petri dish in PBS under a stereoscope for dissection. Fast removed all additional fat and muscles around the eyeball, and another puncture was made in the limbal area to provide easy access for micro-scissors to cut along the limbal ring. Then the cornea, lens, and vitreous, along with the neuroretina, were gently removed. The RPE eyecups were flattened by making four long radial cuts and four additional shorter cuts between them, then transferred to a 48-well plate.

The flatmounts were rinsed twice with PBS, followed by permeabilization with 0.25% PBS-Triton three-time 10min each. Blocking was done for 1hour at room temperature with blocking buffer (1% BSA, 5% DS in 0.25% PBS-T). The flatmounts were incubated with primary antibodies (1:500; V5-Tag, D3H8Q, Cat. #13202, Cell Signaling Technology) overnight at 4 °C. The next day, the flatmounts were washed 3 times with 0.25% PBS-T, 10 min each. And then incubated with secondary antibody (Alexa Fluor® 488, 1:1000 donkey anti-rabbit cat #A21206) in 50% blocking buffer + 50% PBS, 1h at room temperature. Further incubated with Hoechst 33342 (1:2500, Cat. # H1399) and Phalloidin (1:300, Cat. #A12380) for 1h at room temperature, all from Invitrogen. The flatmounts are washed twice for the last time with PBS. Mounted with a drop of Mowiol medium, use a thin No.1 paintbrush to flatten the tissue, and put a coverslip on top of it. After a few hours, seal the coverslip with nail polish.

The slides were stored in the dark at 4°C until imaging using a Zeiss LSM980 confocal scanner with a 10× objective lens for flatmount tiles and a 20× objective lens for the representative area of flatmount. Scans are merged using Zen Blue 3.0 (Zeiss), orthogonal projections, and stitching. Further image processing was performed using Fiji (V2.16.0/1.54p) and Inkscape (V1.2.2)

#### Optical coherence tomography (OCT)

Mice were anesthetized by IP injection of a ketamine/xylazine mixture, and pupils were dilated with one drop of atropine sulfate eye drops (Atropocil®, Edol Pharmaceuticals, Portugal) before imaging. Corneal dehydration and air–corneal interface refraction were prevented using artificial tears and a custom-made polymethylmethacrylate hard contact lens (afocal; curvature, 1.7 mm; diameter, 3.2 mm; Heidelberg Engineering). OCT imaging was performed using a Spectralis OCT system (Heidelberg Engineering) equipped with a +25-diopter add-on lens. Images were acquired with Heidelberg Eye Explorer software (HEYEX, version 2.6.9; Heidelberg Engineering), and the reference pathway length was manually adjusted in the OCT debug window to compensate for the added optics. Confocal scanning laser ophthalmoscopy combined with OCT enabled automatic real-time tracking (ART) and averaging of up to 100 images, reducing speckle noise. After imaging, mice were kept warm, the contact lens was removed and cleaned, and eyes were protected with Corneregel® (Bausch & Lomb) until recovery from anesthesia.

#### Statistical Analysis

Results are presented as Mean ± Standard Error of the Mean (SEM) for independent experiments. Statistical analyses were performed using the GraphPad Prism 9.0.1. software. Normal distribution was assess using the Shapiro-Wilk test. Unpaired student’s t-test and Mann-Whitney test were used to compared between groups if samples followed or not a normal distribution, respectively. Alternatively, one-way ANOVA followed by multiple comparisons Dunnet post hoc correction or one-way ANOVA Kruskal Wallis test were used to compared between groups if samples followed or not a normal distribution, respectively. p-values inferior to 0.05 were considered statistically significant.

## Author Contributions

MCS, ASF and AC conceived and coordinated the study. MLP, ASF, SK, MLS, ECI and LD performed the experiments. MCS, ASF, PA, ST and AC provided support in data analysis and interpretation. MLP wrote the original manuscript draft. MCS, ASF, ST and AC reviewed the manuscript. All authors approved the results and final version of the manuscript.

## Acknowledgments

The authors acknowledge the Cell Tissue Culture, Flow cytometry and Microscopy facilities at NMS, CASO (Centro de Abate de Suínos do Oeste) that generously provided porcine eyes for POS isolations, Boehringer Ingelheim for the gift of AAV-NRF2.

## Data and materials availability

All data needed to evaluate the conclusions in the paper are present in the paper or in the Supplementary Materials.

## Correspondence

and request for materials should be addressed to ASF and MCS.

## Supplementary Figures

**Figure S1.**
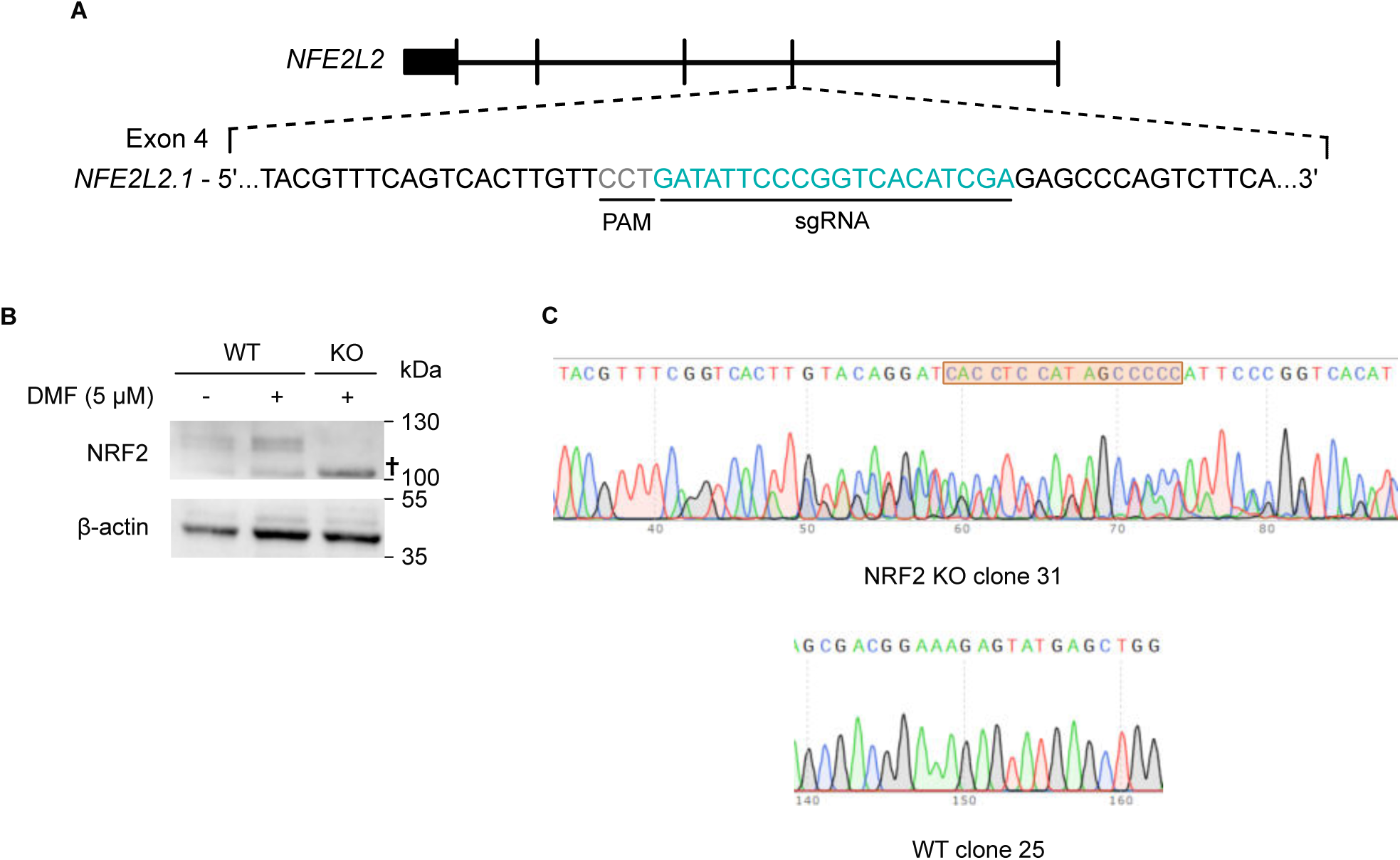
CRISPR/Cas9 strategy to generate ARPE-19 NRF2 KOs. (A) Schematic representation of the strategies designed to generate the NRF2 KO cell line by targeting the *NFE2L2* exon 4. Used sgRNA and Protospacer Adjacent Motif sequences are represented in green and grey, respectively. (B) Western Blot of NRF2 protein levels of NRF2 KO cells upon 5 gM of DMF for 4 hours. (C) Sanger sequencing of the mutation site in NRF2 KO is highlighted.

**Figure S2.**
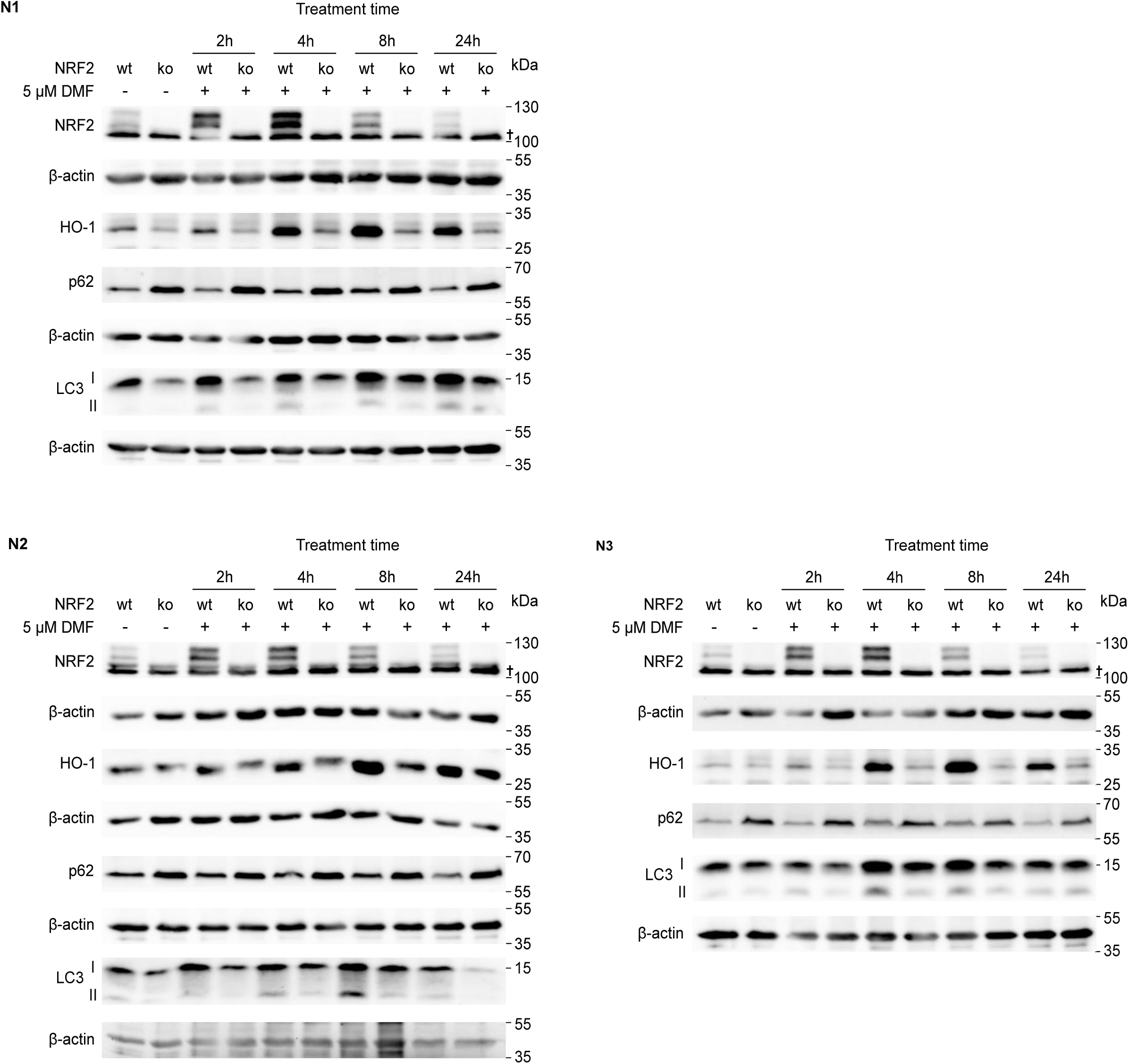
Western Blot of WT and NRF2 KO cells treated for 2, 4, 8 and 24 hours with 5 gM of dimethyl fumarate (DMF), from 3 independent experiments, where the following proteins were accessed: NRF2, HO-1, p62 and LC3. β-actin was used as an endogenous control.

**Figure S3.**
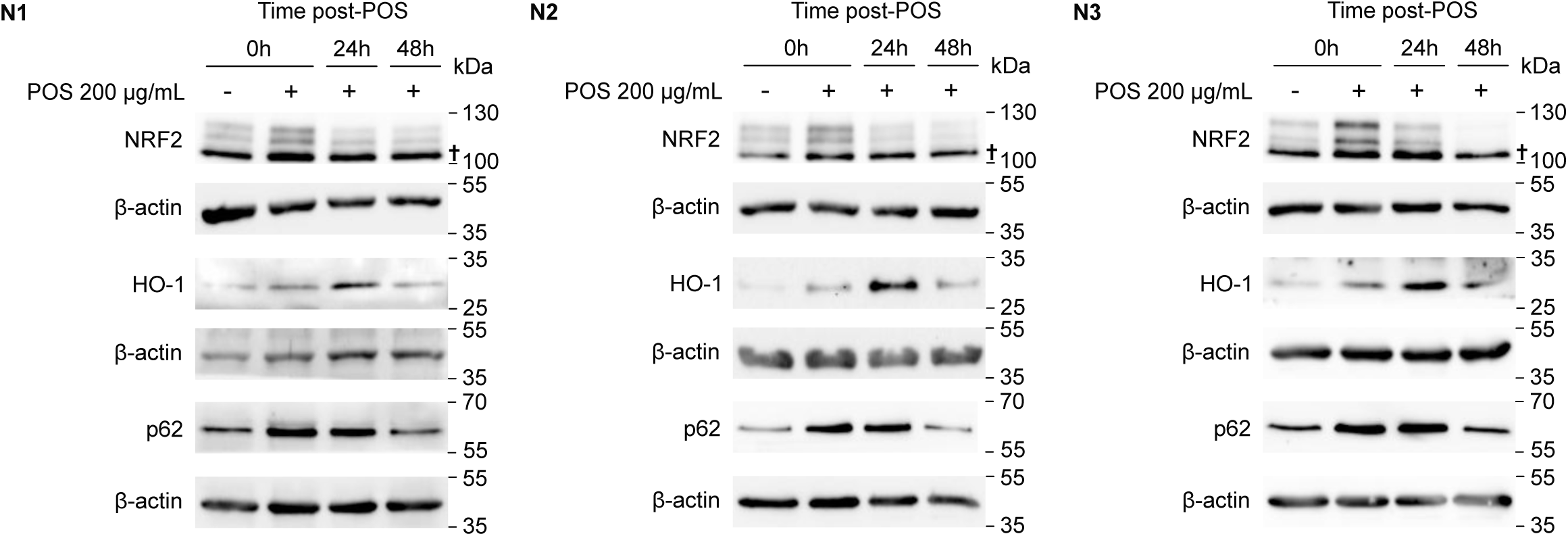
Western Blot of NRF2, HO-1 and p62 protein levels on WT cells incubated 4 hours with POS 200 gg/mL, from 3 independent experiments. β-actin was used as an endogenous control.

**Figure S4.**
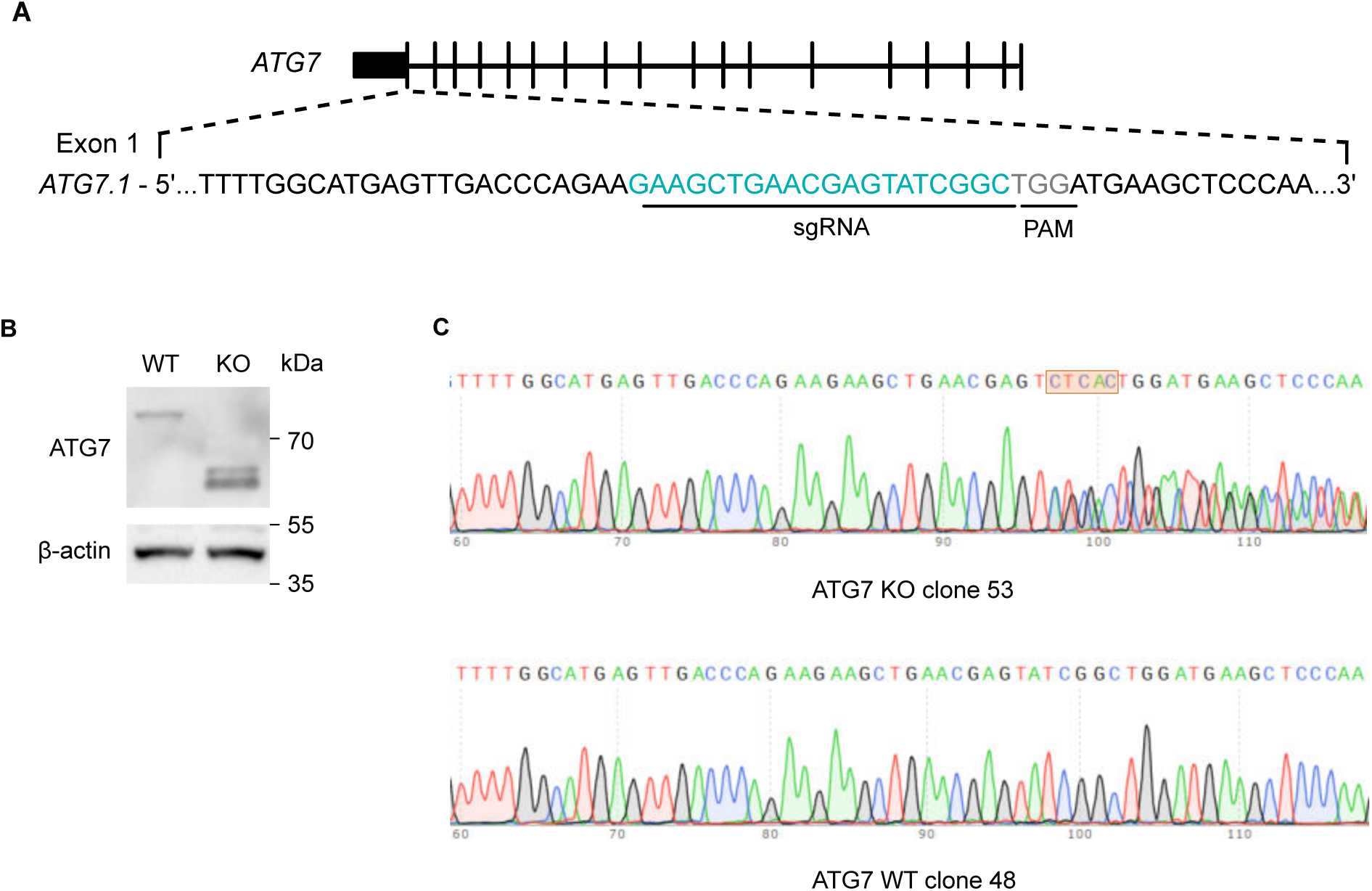
CRISPR/Cas9 strategy to generate ARPE-19 ATG7 KOs. (A) Schematic representation of the strategies designed to generate the ATG7 KO cell line by targeting the *ATG7* exon 1. Used sgRNA and Protospacer Adjacent Motif sequences are represented in green and grey, respectively. (B) Western Blot ATG7 and LC3 protein levels on ATG7 KO cells. (C) Sanger sequencing of the mutation site in ATG7 KO is highlighted.

**Figure S5.**
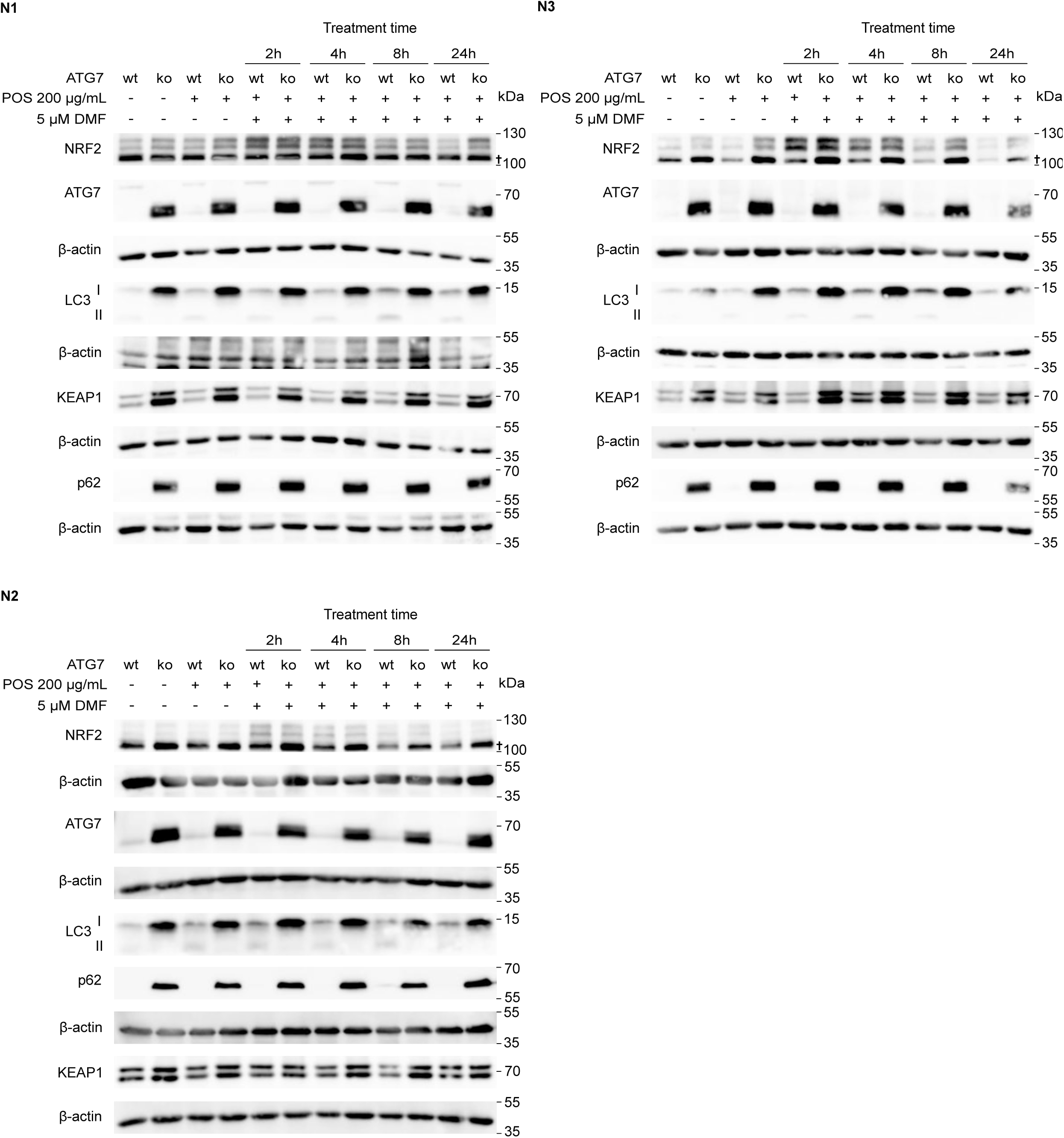
Western Blot of NRF2, ATG7, LC3, KEAP1 and p62 protein levels on WT and ATG7 KO cells incubated 4 hours with POS 200 gg/mL and 48 hours later treated for 2, 4, 8 and 24 hours with 5 gM of dimethyl fumarate (DMF), from 3 independent experiments. β-actin was used as an endogenous control.

**Figure S6.**
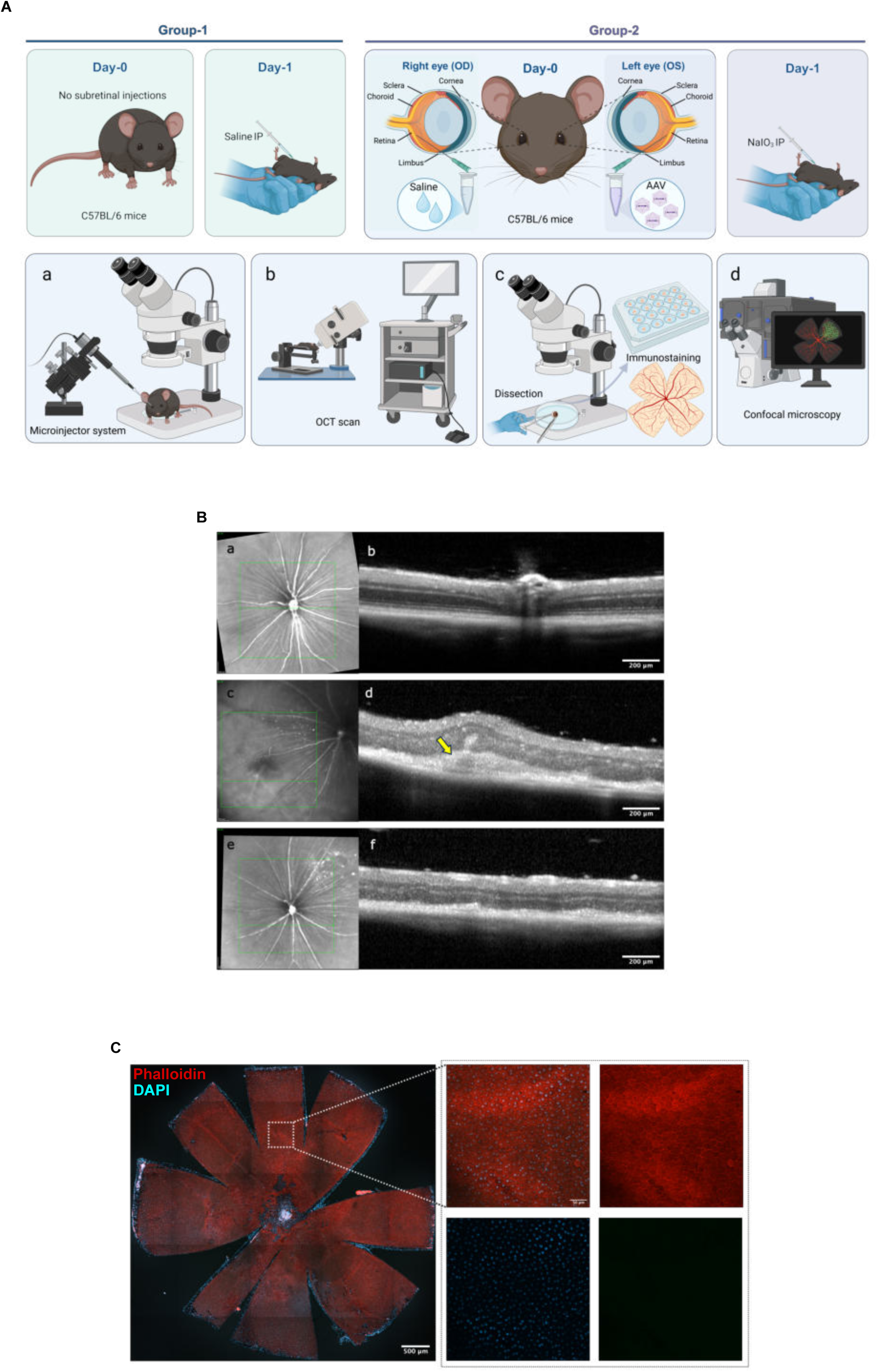
Mice model of RPE degeneration. (A) Experimental design: C57BL/6 mice were divided into 2 groups. Group-1 did not receive subretinal injections, as well as intraperitoneal (IP) injections of NaIO_3_ and Group-2 received subretinal injections at day-0 in the left eyes (AAV-NRF2) and right eyes (saline), using a microinjector (a) and in the next day (day-1), received intraperitoneal (IP) injections of NaIO_3_, to induce oxidative stress in the retina to challenge the RPE cells. Five days after subretinal injection, all eyes were scanned by OCT (b) before sacrificing the mice. After dissection of the eyes, flat mounts were immunostained (c) and images were acquired by confocal microscopy (d). (B) Representative fundus and OCT images of control and NaIO₃-treated mouse eyes with or without AAV-NRF2 subretinal injection. Infrared (IR) fundus images (a, c, e) and corresponding optical coherence tomography (OCT) images (b, d, f) from the same eyes are shown. (a, b) Untreated control mouse eye. (c, d) Mouse eye that received a subretinal AAV-NRF2 injection followed by NaIO₃ administration 1 day later. The arrow indicates a residual subretinal bleb at the injection site. (e, f) The mouse eye that did not receive AAV-NRF2 injection and was administered NaIO₃ on the same day as the mouse shown in panels c and d. Corresponding OCT images illustrate retinal morphology in each experimental condition. (C) Representative immunofluorescence of RPE whole flatmount visualization of the untreated eye (Group 1 mice): Phalloidin (red) staining shows reveals a highly organized, unbroken "honeycomb" mosaic pattern. Scale bar 500 μm (left panel). High-magnification confocal zoom fields (20x magnification, right panels) focusing on the flatmount microstructure: continuous phalloidin staining outlining the characteristic hexagonal pattern of the intact cortical F-actin border. Scale bar 50 μm.

